# A Pairwise Imputation Strategy for Retaining Predictive Features When Combining Multiple Datasets

**DOI:** 10.1101/2022.05.04.490696

**Authors:** Yujie Wu, Boyu Ren, Prasad Patil

## Abstract

In the training of predictive models using high-dimensional genomic data, multiple studies’ worth of data are often combined to increase sample size and improve generalizability. A drawback of this approach is that there may be different sets of features measured in each study due to variations in expression measurement platform or technology. It is often common practice to work only with the intersection of features measured in common across all studies, which results in the blind discarding of potentially useful feature information that is measured only in individual or subsets of all studies. We characterize the loss in predictive performance incurred by using only the intersection of feature information available across all studies when training predictors using gene expression data from microarray and sequencing datasets. We study the properties of linear and polynomial regression for imputing discarded features and demonstrate improvements in the external performance of predictors through simulation and in gene expression data collected on breast cancer patients. We propose and evaluate a pairwise imputation strategy that imputes cross-study missing features in each pair of studies and averages imputed features across pairs. Finally, we provide insights on which subsets of intersected and study-specific features should be used so that missing-feature imputation best promotes cross-study replicability. All code with directions to reproduce results in this paper is available at https://github.com/YujieWuu/Pairwise_imputation

## 1 Introduction

Individual gene expression profiles have successfully been used to model the prognosis or risk of many diseases and disorders in a personalized manner (Van’t Veer *et al*., 2002; Wang *et al*., 2005). These predictive models capture disease identification (Tan and Gilbert, 2003; Fakoor *et al*., 2013), cancer subtyping (Huang *et al*., 2018; Gao *et al*., 2019), and risks of recurrence and relapse (Wang *et al*., 2004; Hartmann *et al*., 2005; Ascierto *et al*., 2012). Technological advancements have seen these studies graduate from custom chips to commercial tools to whole-genome sequencing. This has yielded larger-scale experiments and, over time, the ability to combine multiple gene expression studies of the same disease outcome measured in different patient cohorts. This abundance of data has led to the use of complex statistical prediction and machine learning algorithms for the development of gene signatures (Shipp *et al*., 2002; Ye *et al*., 2003; Pirooznia *et al*., 2008).

A critical issue facing the translation of these gene signatures into viable clinical tests is generalizability, or how well we expect the predictor to perform on a new patient or set of patients. Using techniques such as cross-validation can overestimate how well the predictor will generalize as compared with direct evaluation in a heldout test or validation dataset (Bernau *et al*., 2014). This discrepancy is often due to cross-study heterogeneity in patient characteristics, measurement platforms, and study designs (Patil and Parmigiani, 2018).

To combat the effects of cross-study heterogeneity and increase training sample size with the goal of training a predictor with better generalization properties, researchers have attempted to merge multiple studies (Xu *et al*., 2008). van Vliet *et al*. (2008) showed that pooling data sets together will result in more accurate classification and convergence of signature genes. However, a major challenge in combining these datasets is that the same gene features may not be measured across all studies. This may be due to differences in measurement platform or variations in the same platform when studies are conducted at different points in time.

A common strategy when faced with differing sets of measured genes across studies is to retain only the intersection of gene features found in all studies (Xu *et al*., 2005). We henceforth refer to this method of aggregation as “omitting”, because it simply omits gene expression information that is not measured in at least one study. Taminau *et al*. (2014) proposed a detailed procedure for merging datasets by taking the intersected genes of all studies followed by a batch effect removal procedure. Although omitting provides a simple approach for seamlessly merging studies, it comes with the potentially high cost of discarding important predictive information in features not contained in the intersection. Yasrebi *et al*. (2009) noted that if some genes that have high diagnostic power are not available for all studies, the aggregated data may not actually improve the final predictive model.

A solution to the data loss due to omission is imputation. Zhou *et al*. (2017) built LASSO models to impute missing genes across different studies assayed by two Affymetrix platforms for which the probe names of one platform are a proper subset of the other. Bobak *et al*. (2020) built several imputation models across studies that are measured using a variety of gene expression platforms. Both approaches proceed by first merging the studies together, then using the common genes in the intersection to impute missing genes. As the number of studies increases, the size of the intersection is likely to decrease, resulting in a smaller candidate feature pool and less accurate imputation of omitted genes. This makes merging before imputing a less attractive option when dealing with more than a few studies, such as in the cases of the curated*Data cancer and microbiome dataset collections where dozens of datasets may be available for combination (Gendoo *et al*., 2019; Ganzfried *et al*., 2013). Moreover, these previous approaches were focused on accurate imputation of the missing genes and its effect on downstream analyses such as gene pathway enrichment analysis. Whether or not imputation can help improve a predictive model and make it more generalizable to external data in this context is mostly unstudied.

In this paper, we propose a pairwise imputation procedure, in which instead of merging all available studies together at the outset to build an imputation model for missing gene features, we merge two studies at a time and perform the imputation within the pair. We then repeat this imputation procedure for all possible pairs of studies and average imputed values for features that are missing across multiple pairs. Inspired by the concept of *knowledge transfer* proposed by Vapnik and Izmailov (2015), which posits that some functional forms of the existing features could potentially capture missing information, we examine the ability of both linear and polynomial regression to impute missing features and use LASSO models (Hastie *et al*., 2009) both for imputation and for outcome prediction. Lastly, we consider the impact of using only features selected as “important” (highly associated with the predictive outcome of interest) across both studies within a given study pair (‘Core Imputation’) versus using all available features (‘All Imputation’) when building study-specific imputation models. Here, we revisit the question of whether some form of feature selection and a resulting smaller and more focused set of candidate features is preferable to applying regularization to a larger set of candidate features (Demir-Kavuk *et al*., 2011; Spooner *et al*., 2020).

The paper is organized as follows: Section 2 presents formal notation for the general pairwise imputation framework to impute study-specific missing genes across multiple studies, as well as the specific ‘Core’ and ‘All’ imputation methods. Section 3 presents a simulation study that evaluates the performance of the pairwise imputation method and the ‘Core’ and ‘All’ imputation methods. Section 4 describes a real data analysis predicting the expression of the gene ESR1 across multiple curated breast cancer studies. Section 5 concludes with a discussion.

## 2 Methods

### 2.1 Problem statement

Let *s* = 1, 2, …, *S* index the studies for aggregated analysis, with *n*_*s*_ individuals and *p*_*s*_ genes in study *s*. Let ***X***_*s*_ denote the gene expression data set for the *s*^*th*^ study. ***X***_*s*_ is a *n*_*s*_ *× p*_*s*_ matrix where each row represents an individual and each column represents the measurement values for a particular gene. Let ***Y***_*s*_ be a *n*_*s*_ *×* 1 column vector of the response variable of the *s*-th study. For a pair of studies *s* and *j*, denote by *𝒢*_*sj*_, *𝒢*_*s/j*_ and *𝒢*_*j/s*_ the set of genes that are found in both studies, unique to study *s* and unique to study *j*, respectively. Let |*𝒢*_*sj*_| = *p*_*sj*_, where | *·* | is the cardinality of a set. It follows that |*𝒢*_*s/j*_| = *p*_*s*_ *− p*_*sj*_ and |*𝒢*_*j/s*_| = *p*_*j*_ *− p*_*sj*_. Denote the gene expression matrix for a subset of genes *𝒢* in study *s* as *X*_*s,𝒢*_. Throughout the manuscript, we assume that *p*_*sj*_ *>* 0 for every pair of (*s, j*).

Our goal is to impute the study-specific missing genes to augment the candidate gene set for downstream analysis. To this end, we propose a pairwise imputation approach where imputation is performed among two studies at a time, as the available intersection of genes across any two studies will tend to be larger than that across all *S* studies. For studies *s* and *j*, the pairwise imputation approach uses *𝒢*_*sj*_ to construct prediction models of every gene in *𝒢*_*s/j*_ separately using data from study *s*, based on which the expression profiles in *𝒢*_*s/j*_ will be imputed for study *j*. The same procedure applies to the imputation of genes in *𝒢*_*j/s*_ for study *s*. The imputed studies *s* and *j* both then contain the same set of genes *𝒢*_*s*_ *∪ 𝒢*_*j*_ (see Fig. 1). We repeat this pairwise imputation approach for all 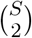 pairs of studies. If a particular gene in one study is imputed multiple times in this process, we will use its averaged measurements over all imputations.

**Figure 1:**
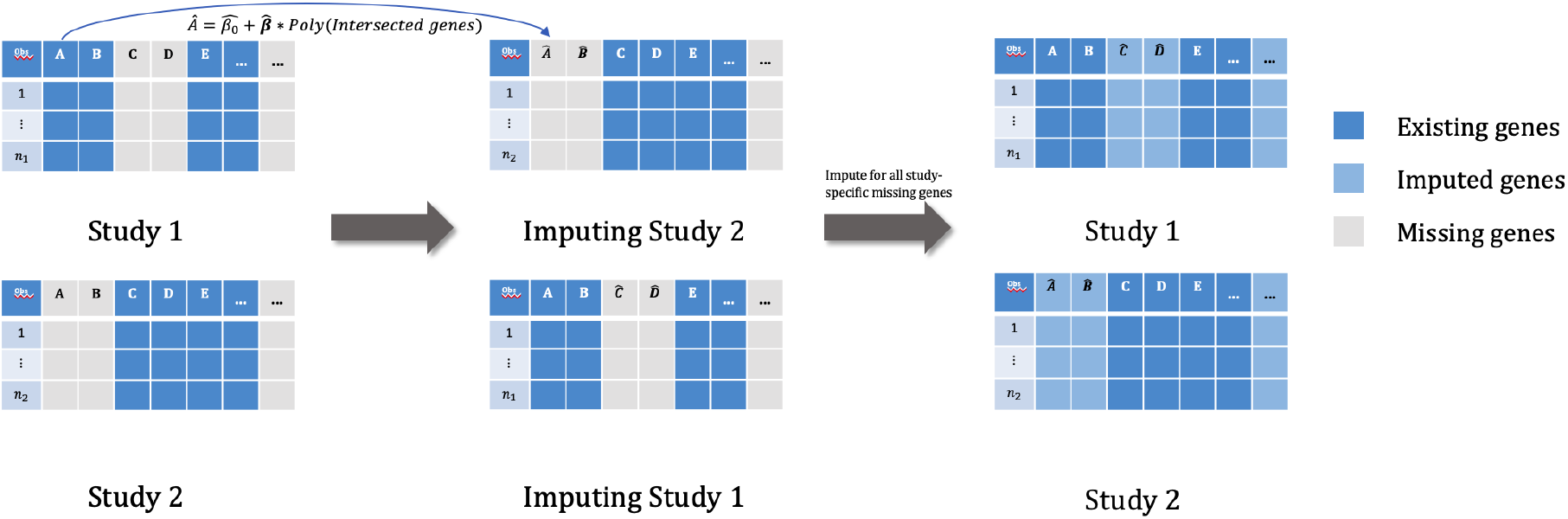
Flow chart of the pairwise imputation approach for study-specific missing genes.

### 2.2 Imputation under incomplete validation set and sparse signals

The pairwise imputation approach has to be refined before it can be used for practical applications. The main limitation of the approach is two-fold: 1) it assumes that all studies are for training and no validation study is present; 2) all genes that appear in at least one study should be included in the final prediction model. The first assumption makes cross-study validation inaccessible while the second can lead to overfitting and increase the computational cost of the approach if *p*_*s*_ is large. Therefore, we propose the following ‘Core’ and ‘All’ variations of the generic pairwise imputation approach to address these two limitations, with ‘Core’ imputation focusing on using selected cross-study genes to build the imputation model and ‘All’ imputation utilizing all available genes in the studies. The algorithm statements of the ‘Core’ and ‘All’ imputation are provided in the supplement.

#### 2.2.1 ‘Core’ imputation

Before introducing the methods, we make the assumption that the response variable is not available in the validation set. For each study, suppose that all genes are not predictive of the outcome, and due to the mixture of signal and noise, a common preprocessing step is to filter to a subset of genes that are most related to the outcome. For example, in each training study we can select the top *q* genes with the largest magnitudes of coefficient estimates from LASSO, where the response is the outcome and the predictors are the expression values of the genes.

The ‘Core’ imputation proceeds by taking two training sets denoted by ***T***_*i*_ and ***T***_*j*_ as a pair, and impute the missing covariates in ***T***_*i*_, ***T***_*j*_ and the validation set (***V***). In the preliminary screening stage, suppose in each training set, the top *q* predictive genes are selected for the final prediction model. Due to study heterogeneity, different sets of genes may be chosen from the two training sets and we denote them as ***Q***_*i*_ and ***Q***_*j*_, respectively. Furthermore, let ***Q***_***V***_ be all the available genes in ***V***, ***ℋ*** = ***Q***_*i*_ *∪* ***Q***_*j*_, ***ℋ***_1_ = ***Q***_*i*_ *∩* ***Q***_*j*_ *∩* ***Q***_***V***_ and ***ℋ***_2_ = ***ℋ****\* ***ℋ***_1_. The idea of ‘Core’ imputation is to impute the genes in ***ℋ***_2_ that are not shared by all of the three studies using genes in ***ℋ***_1_ that are common across all studies. Note that for the ‘Core’ imputation method, the genes used for imputation are all predictive of the outcome in at least one of the training sets. To properly perform ‘Core’ imputation, three different scenarios need to be considered: (1) if a gene is found in only one of ***𝒬***_*i*_ and ***𝒬***_*j*_ and is also missing in ***Q***_***V***_, an imputation model will be built in the training set that has this gene available, and imputation will be performed for the other training set and ***V*** ; (2) if this gene is not missing in ***Q***_***V***_, then no imputation is needed in ***V*** since we can use the original values of this gene; (3) if a gene is available in both ***𝒬***_*i*_ and ***𝒬***_*j*_ but is missing in ***Q***_***V***_, then we merge ***T***_*i*_ and ***T***_*j*_ together to train a single imputation model for this gene and impute in ***V***. If we have *S, S >* 2, training sets, we can repeat the above procedure for all possible 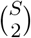 pairs of training sets combined with the additional validation set ***V***, and if a gene is imputed multiple times, we take the average over the multiple imputed values as the final imputation.

We provide a more in-depth illustrative example as well as an algorithm statement in the Supplementary Material.

#### 2.2.2 ‘All’ imputation

The ‘Core’ imputation procedure introduced above will only use the genes in ***ℋ***_1_, which are the genes that are predictive of the outcome, to impute the missing genes in ***ℋ***_2_. However, it is possible that genes not selected for predicting the outcome (i.e. genes not in ***ℋ***) are still helpful for imputing the missing gene expression values. Therefore, another imputation strategy is to use the intersection of all available genes from the three studies instead of focusing only on the intersection of the top predictive genes.

Denote 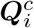 and 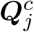 as the other existing genes in ***T***_*i*_ and ***T***_*j*_ but not in ***Q***_***i***_ and ***Q***_***j***_ and let 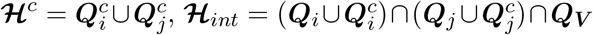. The idea of ‘All’ imputation is to use genes in ***ℋ***_*int*_ to impute the study-specific missing genes in ***ℋ***_2_. Note that the genes in ***ℋ***_*int*_ are the intersection of all the available genes in ***T***_*i*_, ***T***_*j*_ and ***V***, and thus not necessarily predictive of the outcome. Four scenarios require consideration when performing ‘All’ imputation: (1) if a gene is completely missing (e.g. not in ***Q***_*i*_ nor in 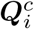) in one of the training sets and ***Q***_***V***_, an imputation model will be built for this gene in the training set that has this gene available and imputation will be performed for the other training set and ***V*** ; (2) if this gene is available in ***Q***_***V***_, then no imputation is needed in ***V*** since we can use the original values of this gene; (3) if the gene is found in ***ℋ***^*c*^ (i.e. this gene is not predictive of the outcome in one of the training sets, but still exists) but is completely missing in ***Q***_***V***_, the training set that has this gene missing in the top *q* predictive gene list can still use its original value, and then we merge the two training sets together to build a single imputation model for this gene and imputation will be performed in ***V*** ; (4) if the gene is found in both ***ℋ***^*c*^ and ***Q***_***V***_, all studies will use their original values and no imputation is needed. An illustrative example and algorithm statement are provided in the Supplementary Material.

## 3 Results

### 3.1 Simulation

#### 3.1.1 Comparison between pairwise and merged imputation methods

We perform a simulation study to compare the performance of our proposed pairwise imputation method to the merged imputation method where we first merge all studies together and use the intersection of variables across all studies to impute study-specific missing variables. We generate four training studies and one external validation study with sample size of 100 for each study, and we evaluate the performance of the imputation methods in terms of the prediction root mean square error (RMSE) in the validation dataset. The overall RMSE is averaged over 300 simulation iterations.

The data for each study is generated from the model,

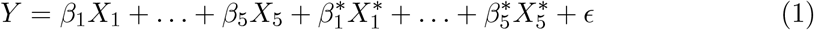

where

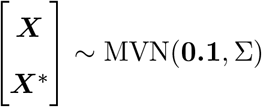

with the variance and covariance being 1 and 0.5, respectively.

To create study-specific patterns of missingness across the four training studies, we fix *X*_1_, *X*_2_ to be common to all studies, while varying the number of missing variables among 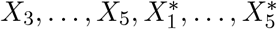 across the four training sets. The validation set is complete, and no imputation is needed.

To predict the outcome of interest, we compare the omitting method where only the intersected variables common across all studies are used for predicting the outcome, pairwise linear and polynomial imputation, and merged linear and polynomial imputation. Table 1 summarizes the imputation models for the study-specific missing variables and the final prediction models for each method.

**Table 1:** Imputation methods considered for comparison

| Method | Imputation Model | Final predicting model |
| --- | --- | --- |
| Omitting | - | LASSO |
| Pairwise linear imputation | LASSO with linear terms of intersected variables | LASSO |
| Pairwise polynomial imputation | LASSO with polynomial terms of intersected variables | LASSO |
| Merged linear imputation | LASSO with linear terms of intersected variables | LASSO |
| Merged polynomial imputation | LASSO with polynomial terms of intersected variables | LASSO |

Figure 2.a shows the RMSE of prediction on the validation set from different imputation methods and the omitting method over the 300 simulation iterations. The omitting method consistently has the worst performance of all methods, and the two pairwise imputation methods have relatively better performance than the corresponding merged imputation methods. To formally compare the performance of different methods by accounting for variation across iterations, we performed a pairwise Wilcoxon tests on the RMSE. Figure 2.b graphically presents the test results, where each method is represented by a single point ordered by the median RMSE over the 300 simulation replicates. The color of the line connecting any two methods indicates the significance level of the test results: a red line indicates that the p-value is less than 0.01; a green line indicates that the p-value is greater than 0.01 but less than 0.05; and a blue line indicates that the p-value is greater than 0.05. For the cases with p-values less than 0.05, we add a directed arrow to indicate the direction of the test, such that the method to which the arrow points has significantly smaller median prediction RMSE. Above each method, we present the proportion of simulation replicates for which that specific method obtained the smallest prediction RMSE in the validation set across the 300 simulation replicates. As shown in the figure, when the number of missing variables is 1 to 4, the pairwise linear imputation method has significantly better performance than the other methods, while when the number of missing variables is 5, 6, and 7, the pairwise polynomial imputation method has slightly smaller prediction RMSE. In Supplementary Figure 1(a), we plot the log RMSE ratio between different imputation methods to the omitting method 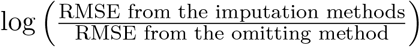 and Supplementary Figure 1(b) shows the log RMSE ratio between the pairwise imputation methods to the merged imputation methods. We also show the average difference in the number of intersected variables used to impute the study-specific missing variables between the merged imputation methods and the pairwise imputation methods as the cross points in Supplementary Figure 1(b). The cross points show that as the number of missing variables increases, the pairwise imputation methods have increasingly larger numbers of intersected variables that can be used to impute the study-specific missing variables as compared to the corresponding merged imputation methods, and the largest discrepancy occurs when the number of missing variables is 3. However, as more variables are missing, the difference approaches 0. This pattern matches with the trend in the log RMSE ratio shown in the same figure, where it initially decreases, but then increases to 0.

**Figure 2:**
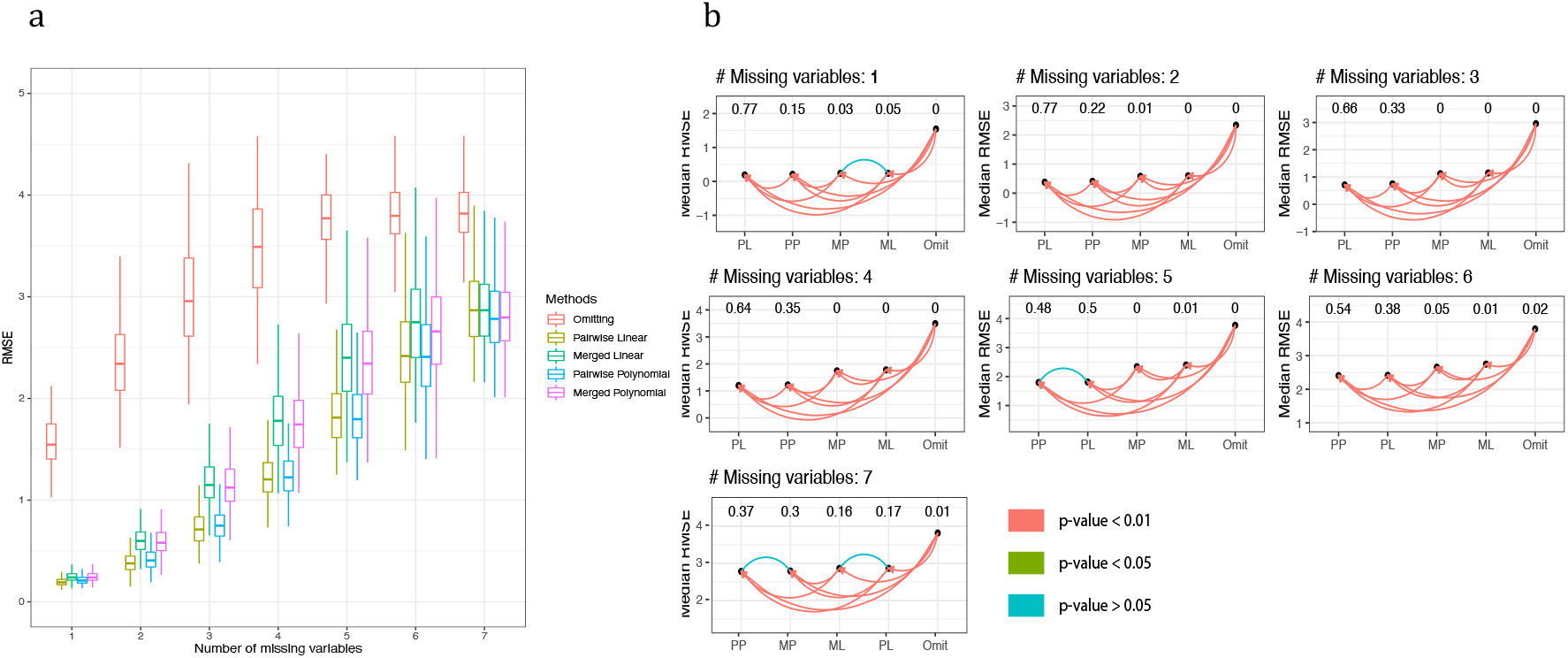
(a) The RMSE of prediction on the validation set for different imputation methods and the omitting method. (b) Pairwise paired Wilcoxon test of the median RMSE of different imputation methods and omitting method over 300 simulation replicates. ‘PP’, ‘MP’, ‘PL’, ‘ML’ stand for pairwise polynomial, merged polynomial, pairwise linear, and merged linear imputation, respectively. A red line indicates that the p-value from the paired Wilcoxon test is less than 0.01; a green line indicates that the p-value is greater than 0.01 but less than 0.05; and a blue line indicates that the p-value is greater than 0.05; and the method to which the arrow is pointing has a significantly smaller median RMSE. The number above each method presents the proportion of times each method has the smallest prediction RMSE in the validation set across the 300 simulation replicates.

Finally, we also consider a scenario that better resembles real gene expression data, where each data set contains both genes that are predictive of the clinical outcome as well as genes that are irrelevant to the outcome. The data generation mechanism is as follows,

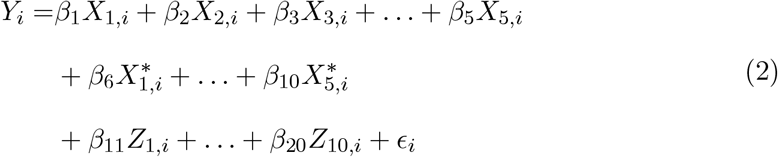

where 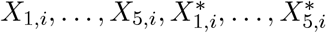, are generated the same way as in Equation (1), while *Z*_1,*i*_, …, *Z*_10,*i*_ follow a multivariate normal distribution with mean 0.1, standard deviation 1 and correlation coefficient 0.2. We restrict the coefficients *β*_11_ = *β*_12_ = … = *β*_20_ = 0, such that *Z*_1_, …, *Z*_10_ can be regarded as the genes that are irrelevant to the clinical outcome. We also vary the number of missing variables in *Z*_1_, …, *Z*_10_ to explore the performance of the pairwise imputation and merged imputation in the presence of missing irrelevant variables. Detailed simulation results can be found in Supplementary Figure 2, where the pairwise imputation method consistently has a smaller RMSE of prediction on the validation set compared to the merged imputation method regardless of the number of missing relevant or irrelevant genes.

#### 3.1.2 Comparison between the ‘Core’ and ‘All’ imputation methods

To mimic the genomic data sets, we performed another simulation study where the signature genes for predicting the outcome of interest are sparse in the whole data set. For illustrative purposes, we have two training sets, and we will make predictions on another validation set that also has missing genes. Since Section 3.1.1 suggests that the pairwise imputation method generally performs better than the merged imputation method in terms of prediction RMSE, in the subsequent simulation study we provide a focused comparison of the ‘All’ and ‘Core’ imputation methods described in Section 2.2 using the pairwise imputation approach.

We generate data as follows, with the sample size of each study set to be 100:

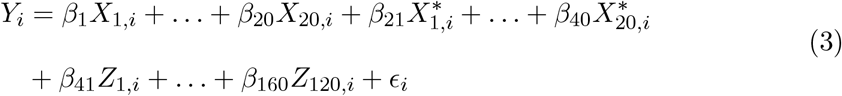

where 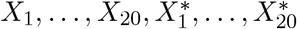 jointly follow a multivariate normal distribution with mean 0.1, variance 1 and covariance 0.5. *Z*_41_, …, *Z*_120_ jointly follows a multivariate normal distribution with mean 0, variance 1 and correlation coefficient 0.2. We restrict the corresponding coefficients *β*_41_, …, *β*_160_ = 0 such that *Z*_1_, …, *Z*_120_ can be regarded as the genes that are irrelevant to the outcome of interest. In simulation, *X*_1_ *− X*_20_ are available to all data sets, and we deliberately set 10 of the genes among 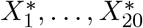, and 50 irrelevant genes among *Z*_1_, …, *Z*_120_ to be missing for both the training and validation sets. Therefore, for each study, we have 20 common predictive genes, 10 study-specific predictive genes and 70 study-specific irrelevant genes.

For the preliminary feature screening, we applied LASSO to select the top *n, n* = 30, 40, …, 100 genes that have largest magnitude of the coefficient in predicting the outcome. Figure 3.a shows the RMSE of prediction on the validation set for the omitting, ‘Core’ and ‘All’ imputation methods. As shown in the figure, both ‘Core’ and ‘All’ imputation methods have better performance than the omitting method, and ‘Core’ imputation consistently has smaller prediction RMSE. Figure 3.b and 3.c show the paired Wilcoxon test results on the RMSE, where the ‘Core’ imputation method has statistically smaller prediction RMSE than the omitting and ‘All’ imputation methods for most of the cases. Supplementary Figure 3(a) shows the boxplots of the log RMSE ratio of the ‘Core’ and ‘All’ imputation methods to the omitting method. Supplementary Figure 3(b) shows the boxplots of the log RMSE ratio of the ‘Core’ imputation method to the ‘All’ imputation method, and the cross points in the same plot show the difference in the average number of irrelevant genes that have non-zero coefficient estimates in the final predictive model of the outcome between the ‘Core’ and ‘All’ imputation. As more irrelevant genes are included, ‘All’ imputation initially has an increasingly large amount of irrelevant genes that have non-zero coefficients in the final predicting model compared to the ‘Core’ imputation method, but the difference gradually approaches 0. Therefore, ‘All’ imputation is more likely than ‘Core’ imputation to have irrelevant genes retained in the final predictive model. Accordingly, the log RMSE ratio between the ‘Core’ and ‘All’ imputation methods initially decreases, and then increases as the difference in the average number of non-zero irrelevant genes in the final predictive model between the two methods gradually approaches 0.

**Figure 3:**
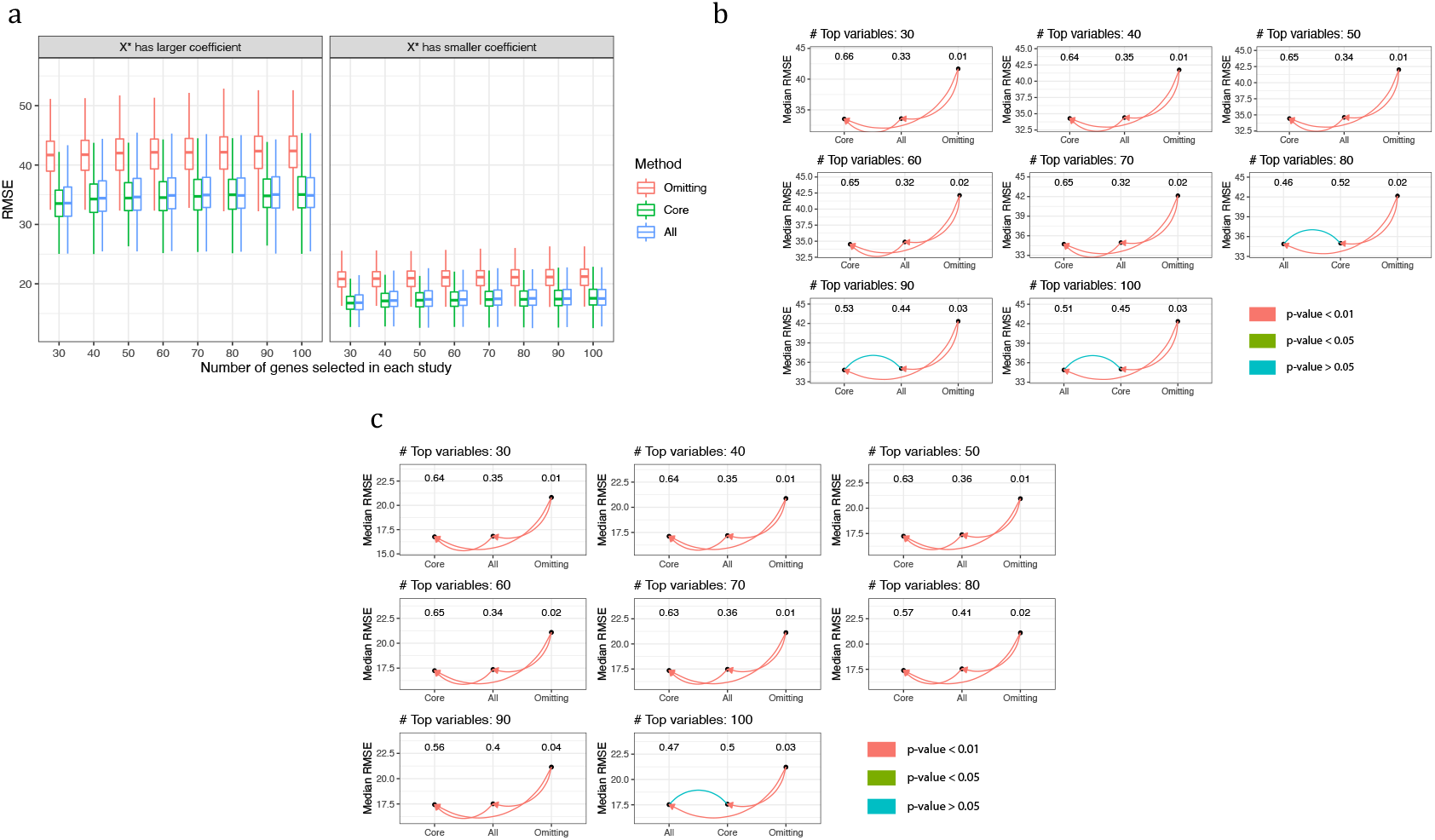
(a) RMSE of prediction on the validation set for the Omitting, ‘Core’ and ‘All’ imputation method across the 300 simulation replicates. Left panel: 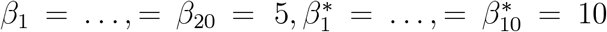 Right panel: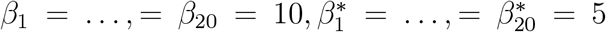; (b, c) Pairwise paired Wilcoxon test on RMSE between Omitting, ‘Core’ and ‘All’ imputation methods for scenarios when *X*^*\**^’s have larger and smaller coefficients than *X*’s, respectively.

In addition, we consider a scenario where *X*_1_, …, *X*_20_ are generated from a multivariate normal distribution but 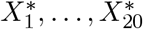 are generated as a complex, non-linear function of *X*_1_, …, *X*_20_, in particular using sine and cosine functions. The simulation results are presented in Supplementary Figures 4 and 5. Contrary to the multivariate normal setting, when *X*^*^,’s are generated as more complex functions of *X*’s, ‘All’ imputation sometimes has smaller prediction error than ‘Core’ imputation, as can be observed for the cases when *X*^*\**^has larger coefficients and top 30 or 40 predictive genes were selected for predicting the outcome. This may be because the preliminary screening process would inevitably select some ‘noise’ features into the top predictive genes while some signals will be neglected, and thus ‘All’ imputation has the advantage of access to more useful genes to impute missing genes compared with the ‘Core’ imputation. Furthermore, *X*^*\**^’s are of complex forms of *X*’s, rendering the imputation to be more difficult as opposed to the multivariate normal case. The gain in the accuracy of the imputed missing genes due to richer information from ‘All’ imputation outweighs the negative effects of more noise genes being included, and therefore yields smaller prediction RMSE in the final predicting model.

### 3.2 Real data analysis

We apply the ‘Core’ and ‘All’ polynomial imputation methods to impute study-specific missing genes on microarray data sets from the ‘curatedBreastData’ Bioconductor package (Planey and Butte, 2013). We selected studies numbered 12093, 16646, 17705, 20181, 20194, 2034, 25055, 25065 because they all used the Affymetrix Human Genome U133A chip for microarray gene expression measurements. The sample sizes of these studies range from 54 to 286, with a total of 1328 patients across studies.

We evaluate the ‘Core’ and ‘All’ polynomial imputation methods to predict the expression level of the gene ESR1. We perform a total of 168 experiments, where for each experiment, two studies are selected as training sets and a third study is chosen as the validation set. The imputation and the final predictive models are all performed using LASSO.

We restrict our analysis to the top 1000 most variable genes in each study. This induces heterogeneous missing patterns across our candidate studies, as the set of 1000 high-variance genes varies across studies. Since not all genes are predictive of the outcome, for the screening step, we fit a LASSO model to predict the ESR1 level based on other gene expression levels in each study and select the genes with larger magnitude of coefficients. We then vary the numbers of top predictive genes we select in each study to predict the expression of ESR1.

Figure 4.a shows the RMSE of prediction on the validation set for the omitting, ‘Core’ and ‘All’ imputation methods. When the number of top predictive genes selected for predicting the outcome in each study is less than 100, both the ‘Core’ and ‘All’ imputation methods have better performance than the omitting method, with ‘All’ imputation having the smallest RMSE. But as the number of predictive genes included in each study exceeds 200, the RMSE from ‘Core’ and ‘All’ imputation methods seem to be similar in performance to omitting. Figure 4.b shows the paired Wilcoxon test results on the RMSE between the three methods. Contrary to the boxplots of the marginal RMSE in Figure 4.a, the ‘Core’ and ‘All’ imputation methods consistently have a significantly smaller prediction RMSE than the omitting method. Thus, 4.b contains the paired information between different methods that could not be reflected by comparing the marginal RMSE of prediction. Supplementary Figure 4a shows the log RMSE ratio of the ‘Core’ and ‘All’ imputation methods to the omitting method as we vary the number of top predictive genes included for predicting the outcome, and Supplementary Figure 4b shows the log RMSE ratio of the ‘Core’ imputation method to the ‘All’ imputation method. Consistent with the paired Wilcoxon test in Figure 4.b, we observe that regardless of the number of genes we have included to predict ESR1 expression levels, the median log RMSE ratios of the ‘All’ and ‘Core’ imputation methods are always smaller than 0, indicating that more than half of the experiments have a decrease in the RMSE by employing the ‘All’ or ‘Core’ imputation method to account for the study-specific missing genes.

**Figure 4:**
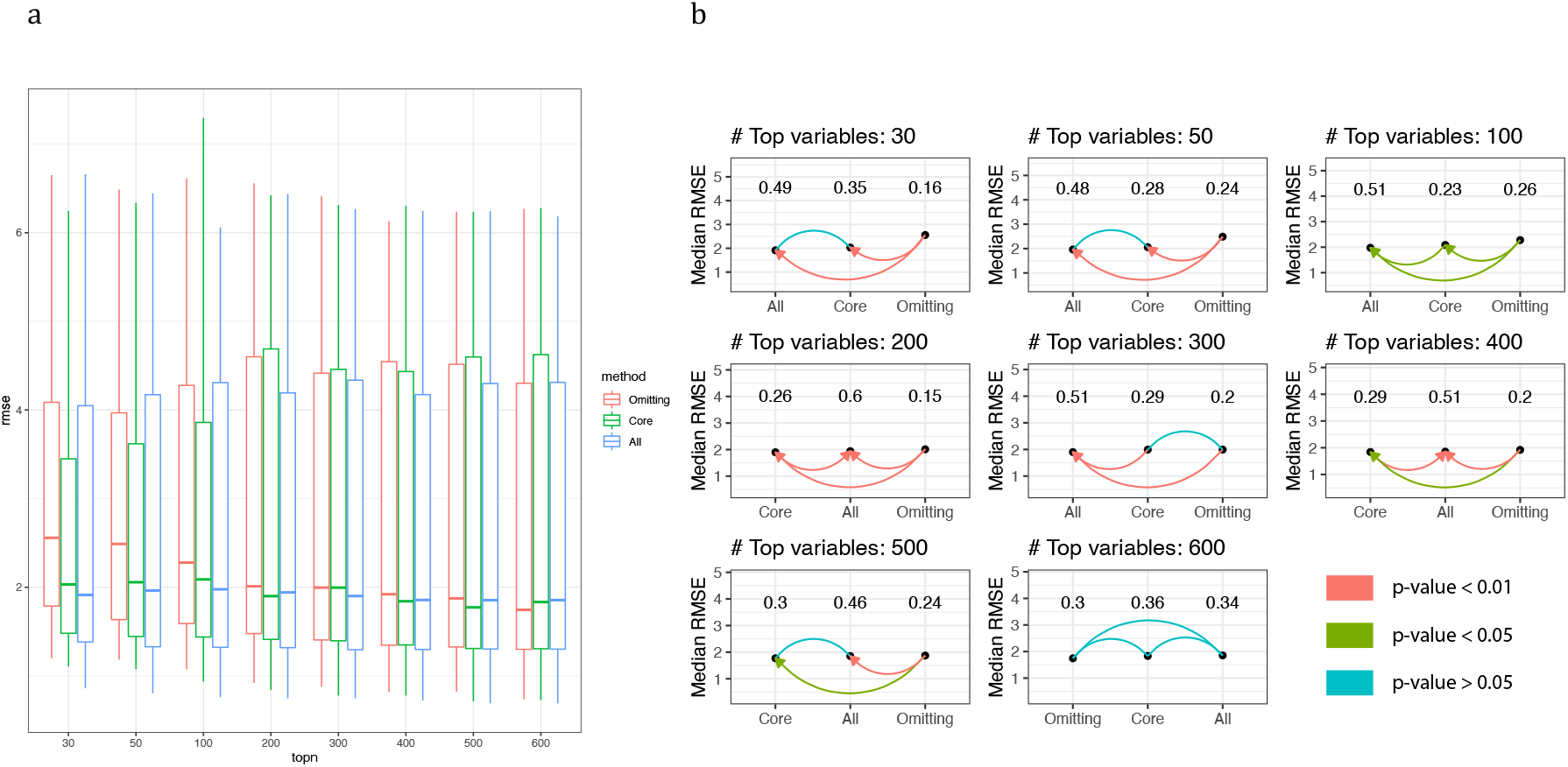
(a) RMSE of prediction on the validation set for Omitting, ‘Core’ and ‘All’ imputation methods. (b) Pairwise paired Wilcoxon test on the RMSE between Omitting, ‘Core’ and ‘All’ imputation methods.

## 4 Discussion

In this paper, we propose a pairwise imputation method to account for differing feature sets across multiple studies when the goal is to combine information across studies to build a predictive model. Compared with the traditionally convenient method of discarding non-intersected genes or the simpler approach of merging studies together and imputing missing genes based on intersected genes shared by all studies, our method has the advantage of utilizing more common genes for imputation based on the intersection between only two studies. Our simulation studies have shown that the pairwise imputation method has significantly better performance than the omitting and merged imputation method in terms of the RMSE of prediction on an external validation set. Moreover, based on the nature of genomics data in which only a subset of genes are likely relevant to the outcome of interest and because the external validation sets may also have genes missing systematically, we compared the ‘Core’ and ‘All’ imputation methods. Our simulation studies here show that both the ‘Core’ and ‘All’ imputation methods will decrease the RMSE of prediction compared to the omitting method, with ‘Core’ imputation demonstrating better performance than the ‘All’ imputation method. Our real data example does not show as strong of a difference between ‘Core’ and ‘All’ imputation. In examining the resulting imputation and prediction models, we find that we are less successful at selecting relevant features using the LASSO feature selection method as we were in the simulation when the data generating mechanism was simple and well-defined. We posit that the advantage of the ‘Core’ imputation method is dependent upon being able to filter an accurate ‘Core’ set of features, and therefore dependent on the quality and performance of the feature selection method used.

When ‘Core’ imputation is successful as in the simulation studies, following Spooner *et al*. (2020) we conjecture that this is because using features that are known to be predictive of the outcome across studies to build imputation models is more reliable and robust to cross-study heterogeneity than using a mixture of cross-study and study-specific features (‘All’ imputation). In ‘All’ imputation, it is possible that a study-specific feature which is only coincidentally predictive within that study will replace a more reliable cross-study feature from the intersection, and while the resulting imputation model would exhibit good performance for that study, it may not generalize well to imputing the same missing feature across studies. We echo the conclusion of Spooner *et al*. (2020) that in some cases, feature selection is more effective than penalization/regularization, and that even if the cross-study selected features are a subset of all features fed to the regularizing model, there may be study-specific features that the regularization prefers.

One limitation of the ‘Core’ and ‘All’ imputation methods is that neither method is using an optimal gene set to impute the study-specific missing genes. ‘Core’ imputation relies only on genes that are predictive of the outcome while completely neglecting other genes that might be informative of those missing genes even though they are not predictive of the outcome. ‘All’ imputation, on other other hand, uses as many genes as possible for imputation with many ‘noise’ genes being included; those additional ‘noise’ features will also lead to less precise imputation of the missing genes. Moreover, an implicit assumption of our imputation procedure is that the genes are missing at random (MAR). If the MAR assumption is violated, for instance, if the missingness mechanism also depends on the outcome, then the imputation might yield even worse predictive performance. We also conduct the bulk of our simulations with a linear model data generating mechanism, which preserves interpretability of the induced missingness patterns, but is likely a simplification of practical data generating mechanisms (however, we do begin to explore more complex associations in the supplement).

Throughout the paper, we applied the linear and polynomial imputation methods, which in essence is training a linear regression of the study-specific missing genes against the linear and polynomial terms of the genes shared by two studies. It would be interesting to employ other complex machine learning techniques for the imputation model, such as Support Vector Machine, Random Forest, Neural Networks to see the relative performance of the pairwise imputation method as well as the ‘ Core’ and ‘All’ imputation methods.

An implicit assumption underlying our method is that there is no or a small amount of heterogeneity across multiple studies. A violation of the homogeneity assumption may lead to the failure of the imputation models. An interesting future research direction is to consider study heterogeneity when imputing the study-specific missing genes.

Moreover, to formally compare the performance of different imputation methods, we applied a preliminary pairwise paired Wilcoxon test on the RMSE of prediction between different methods. Another interesting research topic is on hypothesis tests for rigorous comparison of the performance of different methods that accounts for simulation variation, multiple comparisons and study heterogeneity across multiple studies.

