## supplementary material for "A Pairwise Imputation Strategy for Retaining Predictive Features When Combining Multiple Datasets"

### 1 Notation table

| Notations | Descriptions |
|---|---|
| $s$ | study index |
| $\boldsymbol{X}_s$ | gene profiling dataset for study s |
| $n_s$ | number of observations in study $s$ |
| $p_s$ | number of genes in study $s$ |
| $p_{sj}$ | number of genes common in study $s$ and study $j$ |
| $\boldsymbol{X}_{s,C_{s1}}$ | subset of $\boldsymbol{X}_s$, containing genes that are common across all studies |
| $\boldsymbol{X}_{s,C_{s2}}$ | subset of $\boldsymbol{X}_s$, containing genes that are specific to study $s$ |
| $\mathcal{G}_{sj}$ | genes that are common across study $s$ and study $j$ |
| $\mathcal{G}_{s/sj}$ | genes that are specific to study $s$, but missing in study $j$ |
| $\boldsymbol{T}_1$, $\boldsymbol{T}_2$ | training sets 1 and 2 |
| $\boldsymbol{V}$ | External validation set |
| $\mathcal{H}_1$ | genes that are common in $\boldsymbol{V}$ and the top $n$ genes among $\boldsymbol{T}_1, \boldsymbol{T}_2$ |
| $\mathcal{H}_2$ | union of genes in the top $n$ genes in $\boldsymbol{T}_1$ and $\boldsymbol{T}_2$, but not in $\mathcal{H}_1$ |
| $\mathcal{H}$ | union of the top $n$ genes in $\boldsymbol{T}_1$ and $\boldsymbol{T}_2$ |
| $\mathcal{H}^c$ | union of the remaining genes (i.e., not in the top $n$ genes) in $\boldsymbol{T}_1$ and $\boldsymbol{T}_2$ |
| $\mathcal{H}_{int}$ | intersection of all genes across $\boldsymbol{T}_1, \boldsymbol{T}_2$ and $\boldsymbol{V}$ |

Table 1: Notations

### 2 'Core' and 'All' imputation

#### 2.1 'Core' imputation example

We illustrate the 'Core' imputation on three studies, in which there are two training sets, $s = 1, 2$ and one validation set $s = 3$. Here, we make the assumption that the response variable is not available in the validation set. For each study, suppose there are $p_s$ genes, among which $q_s$ genes are predictive of the outcome (we refer to these genes as signals), and $p_s - q_s$ genes are irrelevant to the outcome (we refer to them as noise). Due to the mixture of signals and noise, a common practice is to filter out genes that are most related to the outcome. Consider, for example, if in each training study we select the top 7 genes with the largest magnitudes of coefficient estimates from LASSO, where the response is the outcome and the predictors are the expression values of the genes. Table 2 provides such an illustrative example, where for notational convenience, we denote the two training sets as $\boldsymbol{T}_1$ and $\boldsymbol{T}_2$ and the validation set as $\boldsymbol{V}$. Let $\mathcal{H}_1 = \{A, B, C\}$ denote the intersected genes among the top 7 genes across the two training sets and all of the available genes in the validation set, $\mathcal{H}_2 = \{D, E, F, G, H, J, K\}$ denote the union of top 7 genes that are not shared by all studies, and $\mathcal{H} = \mathcal{H}_1 \cup \mathcal{H}_2$.

Table 2: Simple example for demonstrating 'Core' Imputation method. The 'Intersected' column contains the intersection of the genes in $\boldsymbol{V}$ and the top 7 genes in $\boldsymbol{T}_1$ and $\boldsymbol{T}_2$; 'Non-intersected' column is for $\boldsymbol{T}_1$ and $\boldsymbol{T}_2$, containing the top 7 genes in $\boldsymbol{T}_1$ and $\boldsymbol{T}_2$ that are not shared by all studies.

| | Top 7 genes for modelling | | Remaining genes |
|---|---|---|---|
| Data set | Intersected | Non-intersected | Not used in 'Core' Imputation |
| $\boldsymbol{T}_1$ | A, B, C | D, E, F, G | ... |
| $\boldsymbol{T}_2$ | A, B, C | G, H, J, K | ... |
| $\boldsymbol{V}$ | A, B, C | E, H, M ,N,... | |

The 'Core' imputation method uses the genes in $\mathcal{H}_1$ to impute the study-specific missing genes in $\mathcal{H}_2$. To perform the 'Core' imputation, three possible scenarios require consideration, and we introduce them using the specific example in Table 2.

1. Gene $D$ in $\mathcal{H}_2$ is found among the top 7 genes of $\boldsymbol{T}_1$, but is missing in $\boldsymbol{V}$ and the top 7 genes in $\boldsymbol{T}_2$. Therefore, we will build an imputation model for gene $D$ using genes in $\mathcal{H}_1$ based on the data from $\boldsymbol{T}_1$, and impute the expression value for gene D in $\boldsymbol{T}_2$ and $\boldsymbol{V}$.

2. Gene $E$ in $\mathcal{H}_2$ is found in $\boldsymbol{V}$ and among the top 7 genes of $\boldsymbol{T}_1$, but is missing in $\boldsymbol{T}_2$. Therefore, the imputation model will be built for gene $E$ using genes in $\mathcal{H}_1$ based on data from $\boldsymbol{T}_1$, and the expression value for gene $E$ will be imputed for $\boldsymbol{T}_2$. Note that even though gene $E$ is also available in the validation set, we will not incorporate it to build the imputation model since $\boldsymbol{V}$ is only used for validation purposes.
3. Gene $G$ in $\mathcal{H}_2$ is found among the top 7 genes of $\boldsymbol{T}_1$ and $\boldsymbol{T}_2$, but is missing in $\boldsymbol{V}$. Therefore, we need to impute gene $G$ for $\boldsymbol{V}$. To fully use the data, we will merge $\boldsymbol{T}_1$ and $\boldsymbol{T}_2$ together and build the imputation model for gene $G$ based on the genes in $\mathcal{H}_1$ from these two training sets.

Genes such as $M$ and $N$ in $\boldsymbol{V}$ are only available in $\boldsymbol{V}$ and are missing in the top 7 genes of $\boldsymbol{T}_1$ and $\boldsymbol{T}_2$. Since $\boldsymbol{V}$ is used for validation, we will not impute Gene $M, N$ for $\boldsymbol{T}_1$ and $\boldsymbol{T}_2$. If we have $S, S > 2$, training sets, we can repeat the above procedure for all possible $\binom{S}{2}$ pairs of training sets combined with the additional validation set $\boldsymbol{V}$, and if a gene is imputed multiple times, we take the average over the multiple imputed values as the final imputation.

#### 2.2 'All' imputation

Denote $\mathcal{H}^c$ as the union of genes that do not belong to the top 7 genes of the two training studies (i.e. the union of genes in the 'Remaining genes' column of Table 3 for $\boldsymbol{T}_1$ and $\boldsymbol{T}_2$), and $\mathcal{H}_{int}$ as the intersected genes of all available genes of the three studies. The 'All' imputation method is to use the genes in $\mathcal{H}_{int}$ to impute the study-specific missing genes in $\mathcal{H}_2$ instead of using only the intersection of the top predictive genes (i.e. genes in $\mathcal{H}_1$). To complete the 'All' imputation, four possible scenarios need considering.

1. Gene $E$ in $\mathcal{H}_2$ is found in $\boldsymbol{V}$ and the top 7 genes of $\boldsymbol{T}_1$, but is missing in $\boldsymbol{T}_2$. Therefore, the imputation model will be built for gene $E$ using genes in $\mathcal{H}_{int}$ based on data from $\boldsymbol{T}_1$, and the expression value of gene $E$ will be imputed for $\boldsymbol{T}_2$. Note that even though gene $E$ is also available in the validation set, we will not incorporate it to build the imputation model since $\boldsymbol{V}$ is only used for validation purposes.
2. Gene $J$ in $\mathcal{H}_2$ is found in $\boldsymbol{V}$ and the top 7 genes of $\boldsymbol{T}_2$, but not in the top 7 genes of $\boldsymbol{T}_1$. However, it is available in $\mathcal{H}^c$ for $\boldsymbol{T}_1$. Therefore, we can directly use the expression value of gene $J$ for $\boldsymbol{T}_1$, and no imputation is needed in this case.
3. Gene $K$ in $\mathcal{H}_2$ is found among the top 7 genes of $\boldsymbol{T}_2$, but not in $\boldsymbol{V}$ and the top 7 genes of $\boldsymbol{T}_1$. However, it is available in $\mathcal{H}^c$ for $\boldsymbol{T}_1$. Therefore, we can directly use the expression value of gene $K$ for $\boldsymbol{T}_1$. Since gene $K$ is completely missing in $\boldsymbol{V}$, we will merge $\boldsymbol{T}_1$ and $\boldsymbol{T}_2$ together and build the imputation model for gene $K$ based on the genes in $\mathcal{H}_{int}$, and then impute the missing gene $K$ value in $\boldsymbol{V}$.
4. Gene $D$ in $\mathcal{H}_2$ is found in $\boldsymbol{T}_1$, but is completely missing in $\boldsymbol{T}_2$ and $\boldsymbol{V}$. Therefore, we will build an imputation model for gene $D$ using genes in $\mathcal{H}_{int}$ based on data from $\boldsymbol{T}_1$, and impute the missing gene $D$ value in $\boldsymbol{T}_2$ and $\boldsymbol{V}$.

Table 3: Simple example for demonstrating 'All' Imputation method. The 'Intersected' column contains the intersection of the genes in $\boldsymbol{V}$ and the top 7 genes in $\boldsymbol{T}_1$ and $\boldsymbol{T}_2$; 'Non-intersected' column is for $\boldsymbol{T}_1$ and $\boldsymbol{T}_2$, containing the top 7 genes in $\boldsymbol{T}_1$ and $\boldsymbol{T}_2$ that are not shared by all studies; 'Remaining genes' column contains the other genes in $\boldsymbol{T}_1$ and $\boldsymbol{T}_2$ that are not in the top 7 gene list.

| | Top 7 genes for modelling | | Remaining genes |
|---|---|---|---|
| Data set | Intersected | Non-intersected | - |
| $\boldsymbol{T}_1$ | A, B, C | D, E, F, G | J, K... |
| $\boldsymbol{T}_2$ | A, B, C | G, H, J, K | ... |
| $\boldsymbol{V}$ | A, B, C | E, H, M ,N, J, ... | |

#### 2.3 Algorithm statement

**Algorithm 1:** 'Core' Imputation

**Data:** $\mathbf{T}_i, \mathbf{T}_j$ (training sets); $\mathbf{V}$ (Validation set)
**Result:** $\widehat{\mathbf{T}}_i, \widehat{\mathbf{T}}_j, \widehat{\boldsymbol{V}}$ (Imputed training and validation sets)
**Initialization**: $\mathcal{Q}_{\boldsymbol{V}}$: All the genes in $\boldsymbol{V}$; select top $q$ predictive genes in each training study: $\mathcal{Q}_i, \mathcal{Q}_j$. Let $\mathcal{H} = \mathcal{Q}_i \cup \mathcal{Q}_j$; $\mathcal{H}_1 = \mathcal{Q}_i \cap \mathcal{Q}_j \cap \mathcal{Q}_{\boldsymbol{V}}$; $\mathcal{H}_2 = \mathcal{H} \backslash \mathcal{H}_1$ ;
**for** *each* $\texttt{gene}_k$ *in* $\mathcal{H}_2$ **do**
    **if** $\texttt{gene}_k \notin \mathcal{Q}_i \cap \mathcal{Q}_j$, *and* $\texttt{gene}_k \notin \mathcal{Q}_{\boldsymbol{V}}$ **then**
        Imputation model: $\texttt{lm(gene}_k \sim \texttt{genes} \in \mathcal{H}_1, \texttt{data} = \texttt{T}_i\texttt{I(gene}_k \in \mathcal{Q}_i) + \texttt{T}_j\texttt{I(gene}_k \in \mathcal{Q}_j\texttt{))}$, and impute for the other two studies
    **end**
    **if** $\texttt{gene}_k \notin \mathcal{Q}_i \cap \mathcal{Q}_j$, *and* $\texttt{gene}_k \in \mathcal{Q}_{\boldsymbol{V}}$ **then**
        Imputation model: $\texttt{lm(gene}_k \sim \texttt{genes} \in \mathcal{H}_1, \texttt{data} = \texttt{T}_i\texttt{I(gene}_k \in \mathcal{Q}_i) + \texttt{T}_j\texttt{I(gene}_k \in \mathcal{Q}_j\texttt{))}$, and impute for the training set having $\texttt{gene}_k$ missing. Validation set retain the original values of $\texttt{Gene}_k$
    **end**
    **if** $\texttt{gene}_k \in \mathcal{Q}_i \cap \mathcal{Q}_j$ *but* $\texttt{gene}_k \notin \mathcal{Q}_{\boldsymbol{V}}$ **then**
        Imputation model: $\texttt{lm(gene}_k \sim \texttt{genes} \in \mathcal{H}_1, \texttt{data} = \texttt{T}_i + \texttt{T}_j\texttt{)}$, and impute for $\boldsymbol{V}$
    **end**
**end**

**Algorithm 2:** 'All' Imputation

**Data:** $\mathbf{T}_i, \mathbf{T}_j$ (training sets); $\mathbf{V}$ (Validation set)
**Result:** $\widehat{\mathbf{T}}_i, \widehat{\mathbf{T}}_j, \widehat{\boldsymbol{V}}$ (Imputed training and validation sets)
**Initialization**: $\mathcal{Q}_{\boldsymbol{V}}$: All the genes in $\boldsymbol{V}$; select top $q$ predictive genes in each training study: $\mathcal{Q}_i, \mathcal{Q}_j$, and let $\mathcal{Q}_i^c, \mathcal{Q}_j^c$ be the other existing genes of each training study. Let $\mathcal{H} = \mathcal{Q}_i \cup \mathcal{Q}_j$; $\mathcal{H}_1 = \mathcal{Q}_i \cap \mathcal{Q}_j \cap \mathcal{Q}_{\boldsymbol{V}}$; $\mathcal{H}_2 = \mathcal{H} \backslash \mathcal{H}_1$; $\mathcal{H}^c = \mathcal{Q}_i^c \cup \mathcal{Q}_j^c$ and $\mathcal{H}_{int} = (\mathcal{Q}_i \cup \mathcal{Q}_i^c) \cap (\mathcal{Q}_j \cup \mathcal{Q}_j^c) \cap (\mathcal{Q}_{\boldsymbol{V}})$ ;
**for** *each* $\texttt{gene}_k$ *in* $\mathcal{H}_2$ **do**
    **if** $\texttt{gene}_k \notin \mathcal{H}^c$ *and* $\texttt{gene}_k \notin \mathcal{Q}_{\boldsymbol{V}}$ **then**
        Imputation model: $\texttt{lm(gene}_k \sim \texttt{genes} \in \mathcal{H}_{int}, \texttt{data} = \texttt{T}_i\texttt{I(gene}_k \in \mathcal{Q}_i) + \texttt{T}_j\texttt{I(gene}_k \in \mathcal{Q}_j\texttt{))}$, and impute for the other two studies
    **end**
    **if** $\texttt{gene}_k \notin \mathcal{H}^c$ *but* $\texttt{gene}_k \in \mathcal{Q}_{\boldsymbol{V}}$ **then**
        Imputation model: $\texttt{lm(gene}_k \sim \texttt{genes} \in \mathcal{H}_{int}, \texttt{data} = \texttt{T}_i\texttt{I(gene}_k \in \mathcal{Q}_i) + \texttt{T}_j\texttt{I(gene}_k \in \mathcal{Q}_j\texttt{))}$, and impute for the other training set. $\boldsymbol{V}$ retains the original values for $\texttt{gene}_k$
    **end**
    **if** $\texttt{gene}_k \in \mathcal{H}^c$ *but* $\texttt{gene}_k \in \mathcal{Q}_{\boldsymbol{V}}$ **then**
        All studies use the original values of $\texttt{gene}_k$, no imputation needed
    **end**
    **if** $\texttt{gene}_k \in \mathcal{H}^c$ *but* $\texttt{gene}_k \notin \mathcal{Q}_{\boldsymbol{V}}$ **then**
        Imputation model: $\texttt{lm(gene}_k \sim \texttt{genes} \in \mathcal{H}_{int}, \texttt{data} = \texttt{T}_i + \texttt{T}_j\texttt{)}$, and impute for $\boldsymbol{V}$.
    **end**
**end**

**Algorithm 3:** Pairwise Imputation

**Data:** $\mathbf{T}_1, \ldots, \mathbf{T}_S$ (Training sets) and $\boldsymbol{V}$ (Validation set)
**Result:** $\widehat{\mathbf{T}}_1, \ldots, \widehat{\mathbf{T}}_S, \widehat{\mathbf{V}}$ (Imputed datasets)
**Initialization**: Form all possible $\binom{S}{2}$ pairs of training sets $(\boldsymbol{T}_i, \boldsymbol{T}_j), 1 \leq i < j \leq S$
**for** *each pair of training sets* **do**
  Run either the 'Core' or 'All' imputation algorithms.
**end**
For each study, if a gene was imputed multiple times, take the simple average of the imputed gene expression values as the final imputation.

**Algorithm 4:** Merged Imputation

**Data:** $\mathbf{T}_1, \ldots, \mathbf{T}_S$ (Training sets) and $\boldsymbol{V}$ (Validation set)
**Result:** $\widehat{\mathbf{T}}_1, \ldots, \widehat{\mathbf{T}}_S, \widehat{\mathbf{V}}$ (Imputed datasets)
**Initialization**: $\mathcal{Q}_{\boldsymbol{V}}$: All the genes in $\boldsymbol{V}$; select top $q$ predictive genes in each training study: $\mathcal{Q}_1, \ldots, \mathcal{Q}_S$. Let $\mathcal{H}_1' = \mathcal{Q}_1 \cap \ldots \cap \mathcal{Q}_S \cap \mathcal{Q}_{\boldsymbol{V}}$, $\mathcal{H}' = \mathcal{Q}_1 \cup \ldots \cup \mathcal{Q}_S$, and $\mathcal{H}_2' = \mathcal{H}' \backslash \mathcal{H}_1'$;
**for** *each* $\texttt{gene}_k$ *in* $\mathcal{H}_2'$ **do**
  Imputation model: $\texttt{lm(gene}_k \sim \texttt{genes} \in \mathcal{H}_1', \texttt{data} = \texttt{T}_1\texttt{I(gene}_k \in \mathcal{Q}_1) + \ldots + \texttt{T}_S\texttt{I(gene}_k \in \mathcal{Q}_S\texttt{))}$, and make imputations.
**end**

### 3 Supplementary figures for Simulation

#### 3.1 Section 3.1

##### 3.1.1 All variables are predictive outcome

##### 3.1.2 Some variables are irrelevant of the outcome

#### 3.2 Simulation with sparse signals and incomplete testing set

#### 3.3 Additional simulation with sparse signals and incomplete testing set when $X^*$'s were generated as a sine and cosine function of $X$'s

### 4 Real data analysis

### 5 Code availability

Code for implementing the proposed methods is available at: https://github.com/YujieWuu/Pairwise_imputation

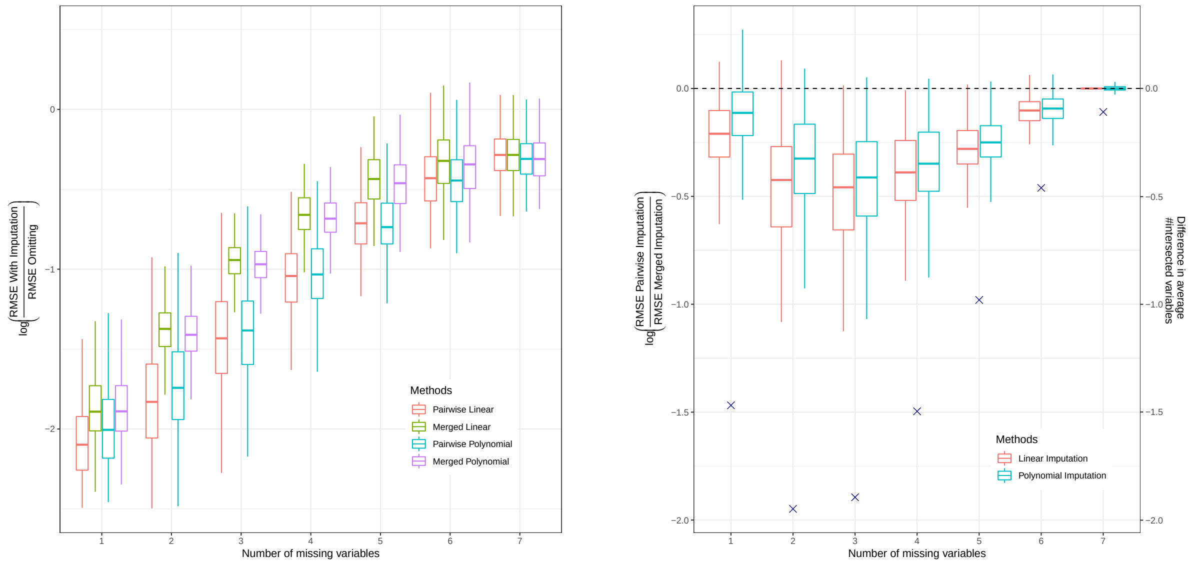


(a) (b)

Figure 1: Log RMSE ratio of prediction RMSE on the validation set. (a) Comparing the RMSE between different merged, pairwise imputation methods with omitting method. (b) Comparing the RMSE between pairwise linear and polynomial imputation models with the corresponding merged imputation methods. The cross points represent the difference in the average number of intersected variables that can be used to impute the study-specific missing variables between the merged and pairwise imputation methods.

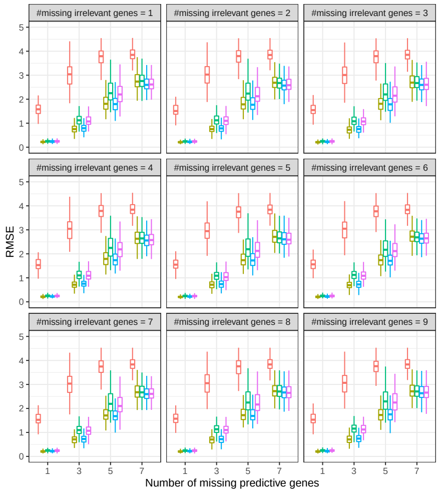


(a)

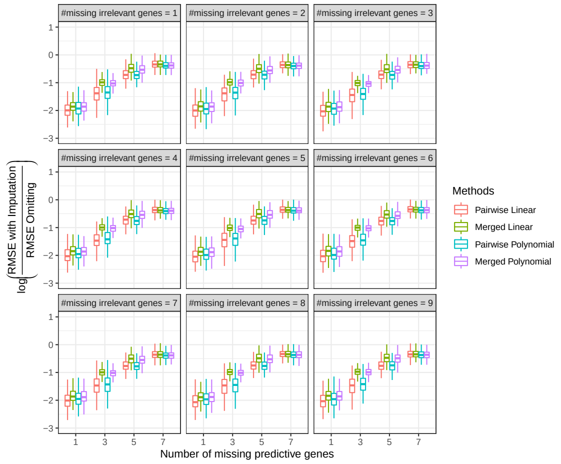


(b)

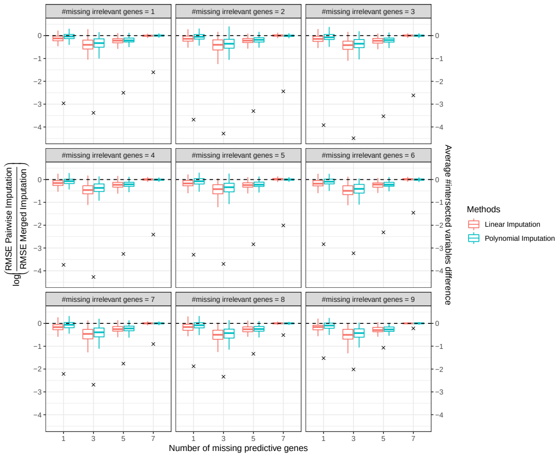


(c)

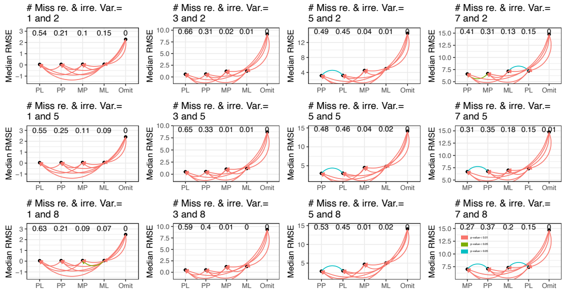


(d)

Figure 2: Simulation results for the scenario when some variables in each study are irrelevant of the outcome. (a) RMSE of prediction on the validation set from different imputation methods. (b) Comparing the RMSE between different merged and pairwise imputation methods with the omitting method. (c) Comparing the RMSE between pairwise linear and polynomial imputation models with corresponding merged imputation methods. The cross points represent the difference in the average number of intersected variables that can be used to impute the study-specific missing variables between the merged and the pairwise imputation models. (d) Wilcoxon paired test on the RMSE of the omitting, pairwise linear, pairwise polynomial, merged linear and polynomial imputation models.

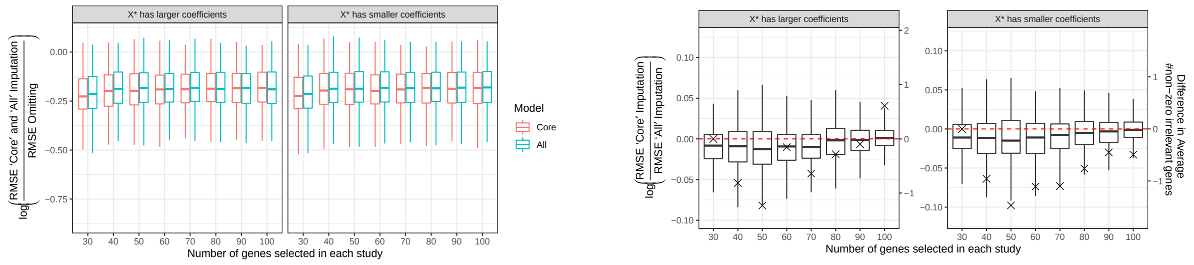


(a) (b)

Figure 3: Simulation results for the scenario when each study has irrelevant genes of the outcome, and the validation set is also incomplete. (a) Comparing the RMSE of prediction on the validation set between 'Core' and 'All' imputation methods with omitting method. (b) Comparing the RMSE from the 'Core' imputation method to the 'All' imputation method. The cross points represent the difference in the average number of irrelevant genes that have non-zero coefficients in the final predicting model between the 'Core' and 'All' imputation methods.

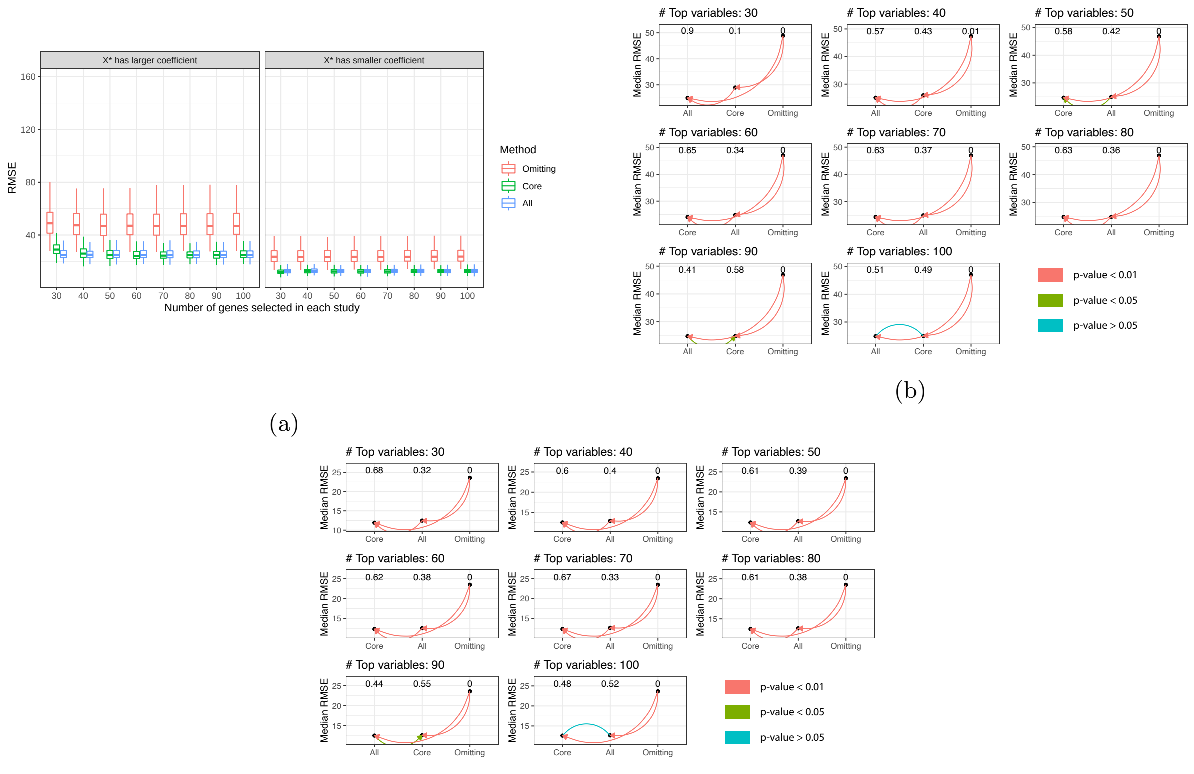


(a) (b) (c)

Figure 4: Simulation results for the scenario when $X^*$'s were generated as sine and cosine functions of $X$'s. (a) RMSE of prediction on the validation set for the Omitting, 'Core' and 'All' imputation method across the 300 simulation replicates. Left panel: $\beta_1 = \ldots, = \beta_{20} = 5, \beta_1^* = \ldots, = \beta_{10}^* = 10$; Right panel: $\beta_1 = \ldots, = \beta_{20} = 10, \beta_1^* = \ldots, = \beta_{20}^* = 5$; (b, c) Pairwise paired Wilcoxon test on RMSE between Omitting, 'Core' and 'All' imputation methods for scenarios when $X^*$'s have larger and smaller coefficients than $X$'s, respectively.

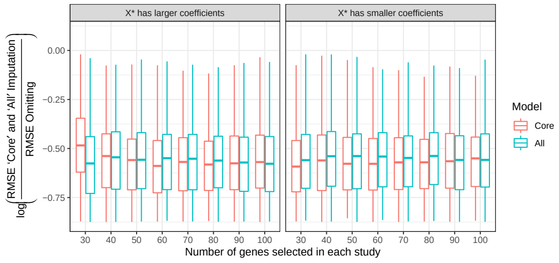


(a)

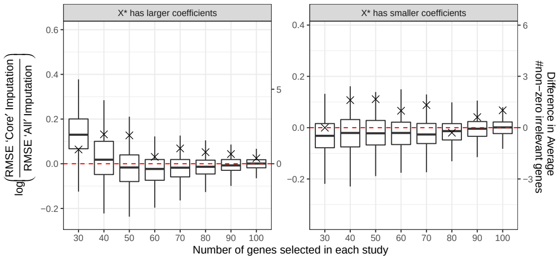


(b)

Figure 5: Simulation results for the scenario when $X^*$'s were generated as sine and cosine functions of $X$'s. (a) Comparing the RMSE of prediction on the validation set between 'Core' and 'All' imputation methods with omitting method. (b) Comparing the RMSE from the 'Core' imputation method to the 'All' imputation method. The cross points represent the difference in the average number of irrelevant genes that have non-zero coefficients in the final predicting model between the 'Core' and 'All' imputation methods.

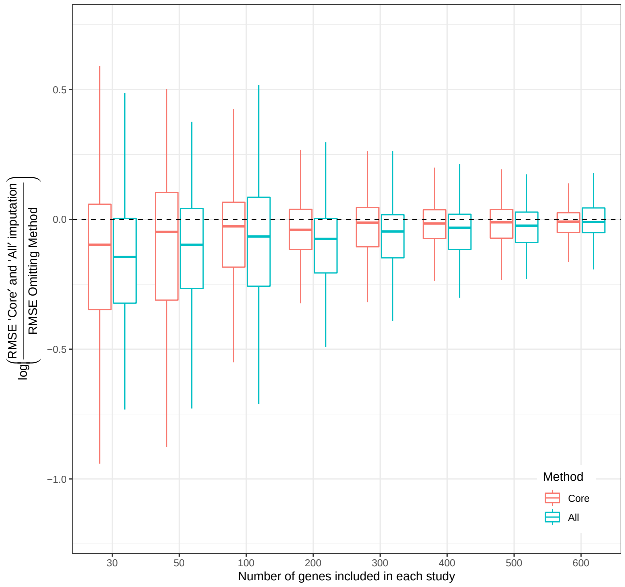


(a)

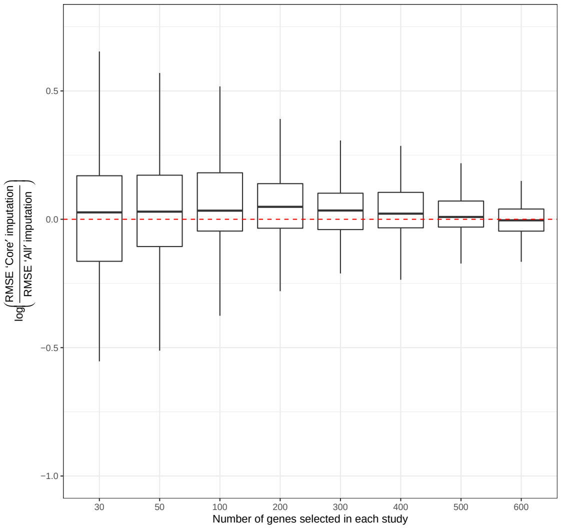


(b)

Figure 6: Results for real data analysis. (a) Comparing RMSE of prediction on the validation study between the 'Core' and 'All' imputation methods to the omitting method. (b) Comparing the RMSE between the 'Core' imputation to the 'All' imputation method.
